# Genome survey of the freshwater mussel *Venustaconcha ellipsiformis* (Bivalvia: Unionida) using a hybrid *de novo* assembly approach

**DOI:** 10.1101/265157

**Authors:** Sébastien Renaut, Davide Guerra, Walter R. Hoeh, Donald T. Stewart, Arthur E. Bogan, Fabrizio Ghiselli, Liliana Milani, Marco Passamonti, Sophie Breton

## Abstract

Freshwater mussels (Bivalvia: Unionida) serve an important role as aquatic ecosystem engineers but are one of the most critically imperilled groups of animals. Here, we used a combination of sequencing strategies to assemble and annotate a draft genome of *Venustaconcha ellipsiformis*, which will serve as a valuable genomic resource given the ecological value and unique “doubly uniparental inheritance” mode of mitochondrial DNA transmission of freshwater mussels. The genome described here was obtained by combining high coverage short reads (65X genome coverage of Illumina paired-end and 11X genome coverage of mate-pairs sequences) with low coverage Pacific Biosciences long reads (0.3X genome coverage). Briefly, the final scaffold assembly accounted for a total size of 1.54Gb (366,926 scaffolds, N50 = 6.5Kb, with 2.3% of “N” nucleotides), representing 86% of the predicted genome size of 1.80Gb, while over one third of the genome (37.5%) consisted of repeated elements and more than 85% of the core eukaryotic genes were recovered. Given the repeated genetic bottlenecks of *V. ellipsiformis* populations as a result of glaciations events, heterozygosity was also found to be remarkably low (0.6%), in contrast to most other sequenced bivalve species. Finally, we reassembled the full mitochondrial genome and found six polymorphic sites with respect to the previously published reference. This resource opens the way to comparative genomics studies to identify genes related to the unique adaptations of freshwater mussels and their distinctive mitochondrial inheritance mechanism.

## Introduction

Through their water filtration action, freshwater mussels (Bivalvia: Unionida) serve important roles as aquatic ecosystem engineers (Gutiérrez et al. 2003; Spooner & Vaughn 2006), and can greatly influence species composition (Aldridge et al. 2007). From a biological standpoint, they are also well known for producing obligate parasitic larvae that metamorphose on freshwater fishes (Lopes-Lima et al. 2014), for being slow-growing and long-lived, with several species reaching >30 years old and some species >100 years old (see Haag & Rypel 2011 for a review), and for exhibiting an unusual system of mitochondrial transmission called Doubly Uniparental Inheritance or DUI (see Breton et al. 2007; Passamonti & Ghiselli 2009; Zouros 2013 for reviews). From an economic perspective, freshwater mussels are also exploited to produce cultured pearls (Haag 2012). Regrettably however, habitat loss and degradation, overexploitation, pollution, loss of fish hosts, introduction of non-native species, and climate change have resulted in massive freshwater mussel decline in the last decades (reviewed in Lopes-Lima et al. 2017; 2018). For example, more than 70% of the ~300 North American species are considered endangered at some level (Lopes-Lima et al. 2017).

While efforts are currently underway to sequence and assemble the genome of several marine bivalves such as the mussel *Mytilus galloprovincialis* (Murgarella et al. 2016), genomic resources for mussels in general are still extremely scarce. In addition to *M. galloprovincialis*, the genomes of several other mytilid mussel species, such as the deep-sea vent/seep mussel *Bathymodiolus platifrons*, the shallow-water mussel *Modiolus philippinarum* and the golden mussel *Limnoperna fortunei* have recently been published (Sun et al. 2017, Uliano-Silva et al. 2018). In all cases, genomes have proven challenging to assemble due to their large size (~1.6 to 2.4Gb), widespread presence of repeated elements (~30% of the genome, and up to 62% of the genome for the shallow-water mussel *M. philippinarum*, Sun et al. 2017) and high heterozygosity (e.g. Murgarella et al. 2016, Mu et al. 2017, Uliano-Silva et al. 2018). For example, the *M. galloprovincialis* genome remains highly fragmented, with only 15% of the gene content estimated to be complete (Murgarella et al. 2016). With respect to freshwater mussels (order Unionida), no nuclear genome draft currently exists. An assembled and annotated genome for freshwater mussels has the potential to be utilized as a valuable resource for many researchers given the biological value and threatened features of these animals. In addition, we predicted that, contrary to most other sequenced bivalve species, heterozygosity of *V. ellipsiformis* would be relatively low, given its history of genetic bottlenecks due to repeated glaciations events over its current geographical distribution (Zanatta & Harris 2013). Genomic resources are needed to help identifying genes essential for survival (and/or the genetic mechanisms that led to decline) and ultimately for developing monitoring tools for endangered biodiversity and plan sustainable recoveries (Pavey et al. 2016; Savolainen et al. 2013). In addition, a sequenced genome will help answer more fundamental questions of sex-determination (Breton et al. 2011; 2017) and genome evolution through comparative genomics approaches (e.g. Sun et al. 2017).

Given the challenges in assembling a reference genome for saltwater mussels (Sun et al. 2017; Murgarella et al. 2016), we used a combination of different sequencing strategies (Illumina paired-end and mate pair libraries, Pacific Biosciences long reads, and a recently assembled reference transcriptome, Capt et al. 2018) to assemble the first genome draft in the family Unionidae. Hybrid sequencing technologies using long read–low coverage and short read–high coverage offer an affordable strategy with the advantage of assembling repeated regions of the genome (for which short reads are ineffective) and circumventing the relatively higher error rate of long reads (Koren et al. 2012; Miller et al. 2017). Here, we present a *de novo* assembly and annotation of the genome of the freshwater mussel *V. ellipsiformis*.

## Methods

To determine the expected sequencing effort to assemble the *V. ellipsiformis* genome, i.e., the necessary software and computing resources required, we first searched for C-values from other related mussel species. C-values indicate the amount of DNA (in picograms) contained within a haploid nucleus and is roughly equivalent to genome size in megabases. Two closely related freshwater mussel species in the same order (Unionida) as *V. ellipsiformis* (*Elliptio* sp., c-value = 3; *Uniomerus* sp., c-value = 3.2), in addition to other bivalve species from different orders (e.g. *Mytilus* spp. [order Mytilida], c-value = 1.3–2.1; *Dreissena polymorpha* [order Venerida], c-value = 1.7) were identified on the Animal Genome Size Database (http://www.genomesize.com). As such, we estimated the *V. ellipsiformis* genome size to be around ~1.5–3.0Gb, and this originally served as a coarse guide to determine the sequencing effort required, given that when the sequencing for *V. ellipsiformis* was originally planned, no mussel genome had yet been published.

### Mussel specimen sampling, genomic DNA extraction and library preparation

Adult specimens of *V. ellipsiformis* were collected from Straight River (Minnesota, USA; Lat 44.006509, Long −93.290899), and species was identified according to Badra (2007). Specimens were sexed by microscopic examination of gonad smears. Gills were dissected from a single female individual and genomic DNA was extracted using a Qiagen DNeasy Blood & Tissue Kit (QIAGEN Inc., Valencia, CA, USA) using the animal tissue protocol. The quality and quantity of DNA, respectively, were assessed by electrophoresis on 1% agarose gel and with a BioDrop mLITE spectrophotometer (a total of 15 µg of DNA was quantified using the spectrophotometer). For whole genome shotgun sequencing and draft genome assembly, we used two sequencing platforms: Illumina (San Diego, CA) Hiseq2000 and Pacific Biosciences (Menlo Park, CA) PacBio RSII. First, three paired-end libraries with insert size of 300bp were constructed using Illumina TruSeq DNA Sample Prep Kit. One mate pair library with insert sizes of about 5Kb was constructed for scaffolding process using Illumina Nextera mate-pair library construction protocol. For high-quality genome assembly, Pacific Biosciences system was employed for final scaffolding process using long reads. Pacific Biosciences long reads (>10Kb) were generated using SMRT bell library preparation protocol (ten SMRT cells were sequenced). Construction of sequencing libraries and sequencing analyses were performed at the Genome Quebec Innovation Centre (McGill University, Qc, Canada).

### Pre-processing of sequencing reads

We quality trimmed paired-end and mate-pair reads using Trimmomatic 0.32 (Bolger et al. 2014) with the options illuminaclip:TruSeq3-PE.fa:2:30:10 leading:3 trailing:3 slidingwindow:6:10 minlen:36. This allowed removal of base pairs below a threshold Phred score of three at the leading and trailing end, in addition to removing base pairs based on a sliding window calculation of quality (mininum Phred score of ten over six base pairs). Finally, if trimmed reads fell below a threshold length (36bp), both sequencing pairs were removed. We verified visually the quality (including contamination with Illumina paired-end adaptors) before and after trimming using fastqc (Andrews 2010). This allowed us to only keep high quality reads prior to the assembly steps.

Following quality trimming, we used bfc (Li & Durbin 2009) to perform error correction for the Illumina paired-end sequencing data. bfc suppresses systematic sequencing errors, which helps to improve the base accuracy of the assembly and reduce the complexity of the *de Bruijn* graph based assembly, described below.

Corrected paired-end reads were subsequently used to identify the optimal K value that provides the most distinct genomic k-mers using KmerGenie v1.7016 (Chikhi & Medvedev 2014). We tested k = 10 to 100, in incremental steps of 10, and we then refined the interval from 20 to 40, in incremental steps of 2 to get a more precise estimate of K.

### Genome size and heterozygosity estimation

We used jellyfish 2.1.4 (Marcais et al. 2011) for counting k-mers of lengths 17, 19, 21, 31 and 41, and obtain their frequency distributions, using the error-corrected, trimmed paired end reads. Based on k-mer frequency distributions, we then used genomescope (Vurture et al. 2017) to estimate the overall characteristics of the genome, including genome size, heterozygosity rate and repeat content in R version 3.4.4 (R Core Team 2012). GenomeScope attempts to fit mixture models of four evenly spaced negative binomial distributions to each k-mer profile in order to measure the relative abundances of heterozygous, homozygous, unique and duplicated sequences.

### Genome assembly strategy

We used ABySS 2.0 (Jackman et al. 2017), a modern genome assembler specifically built for large genomes and reads acquired by different sequencing strategies. ABySS 2.0 works similarly to ABySS (Simpson et al. 2009), by using a distributed *de Bruijn* graph representation of the genome, therefore allowing parallel computation of the assembly algorithm across a network of computers. In addition, the software makes use of long sequencing reads (Illumina mate-pair libraries and Pacific BioSciences long reads) to bridge gaps and scaffold contigs. Yet, as memory requirements and computing time scale up exponentially with genome size, for large genomes (>1Gb), these rapidly become very large (>100GB of RAM) and unpractical. Consequently, Jackman et al. (2017) introduced ABySS 2.0, which employs a probabilistic data structure called a Bloom filter (Bloom 1970) to store a *de Bruijn* graph representation of the genome and, consequently, greatly reduces memory requirements and computing time. The Bloom filter allows removing from memory the majority of nearly identical k-mers likely caused by sequencing errors, as k-mers with an occurrence count below a user-specified threshold are discarded. The caveat is that it can generate false positive extension of contigs, but through optimization, this can be kept well below 5%, and in fact, false positives can be corrected later on in the assembly step (Jackman et al. 2017).

In the current study, we combined different types of high throughput sequencing to aid in assembling the genome (**Table 1**). ABySS 2.0 (Jackman et al. 2017) performs a first genome assembly step without using the paired-end information, by extending unitigs until either they cannot be unambiguously extended or come to an end due to a lack of coverage (*uncorrected unitigs*). This first *de Bruijn* graph representation of the genome is further cleaned of vertices and edges created by sequencing errors (*unitigs*). Paired-end information is then used to resolve ambiguities and merge *contigs*. Following this, mate-pairs are mapped onto the assembly to create *scaffolds*, and finally long reads (Pacific Biosciences long reads) and the *V. ellipsiformis* reference transcriptome from Capt et al. (2018) were also mapped onto the assembly to create *long-scaffolds*. This reference transcriptome was assembled from a pool of sequences coming from four different male and female individuals and further details are provided in Capt et al. (2018). Although ideally sequencing information would all come from a single individual, the current study design did not allow for this. In addition, given that coding sequences are conserved compared to non-coding regions, it remains highly valuable to use a transcriptome in a *de novo* genome assembly.

**Table 1:**
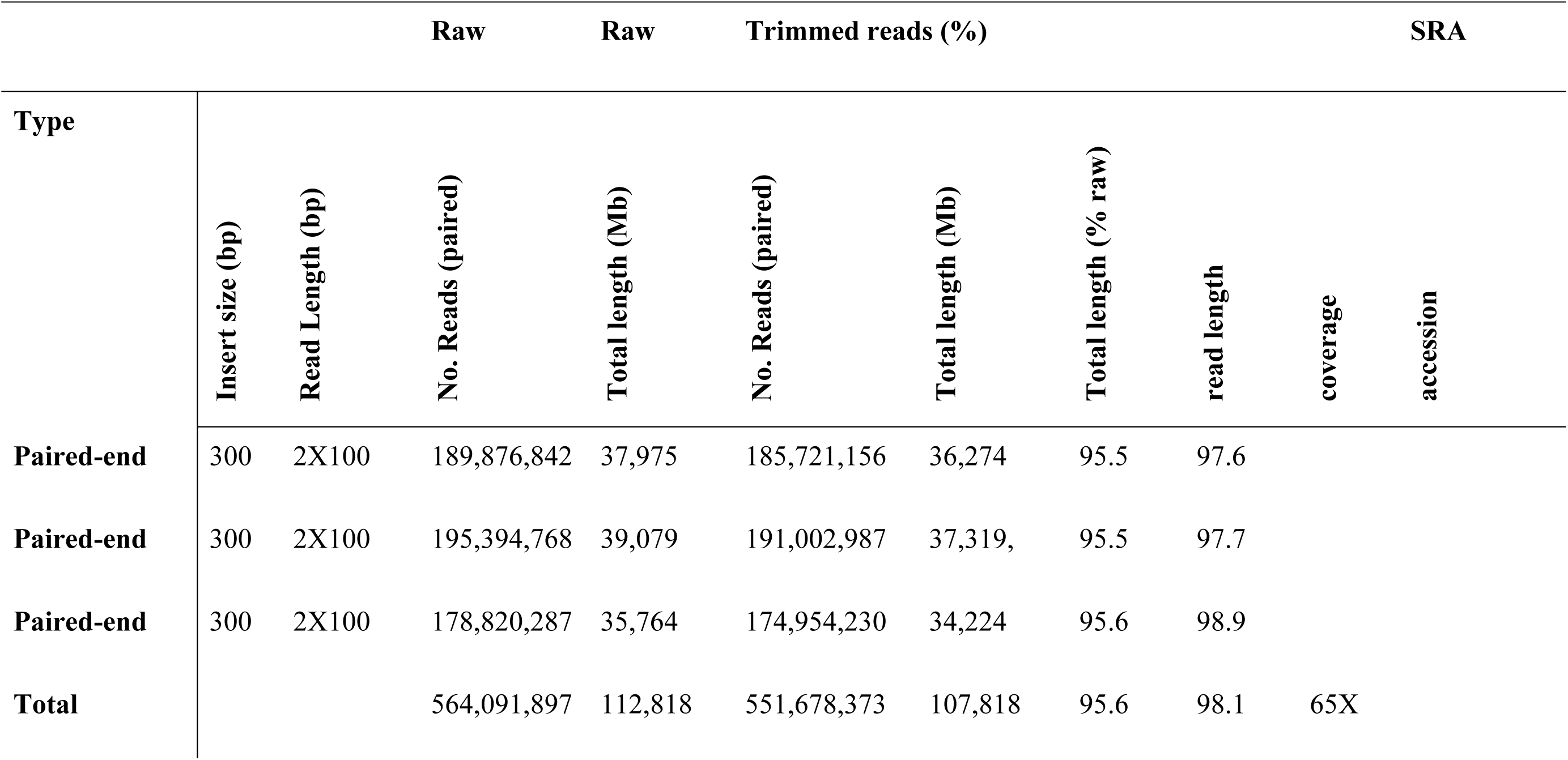
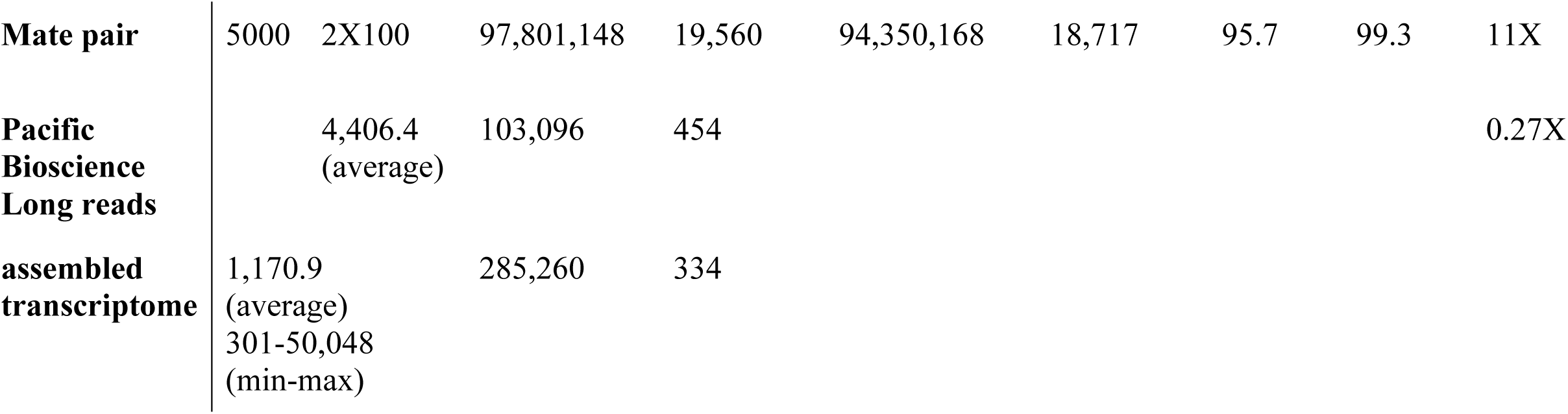
DNA sequencing strategy.

We ran the ABySS 2.0 assembly stage (abyss-bloom-dbg) with a k-mer size of 41 (ABySS requires an odd number k-mer), a Bloom filter size of 24GB, 4 hash functions and a threshold of k-mer occurrence set at 3. These parameters were chosen after performing several test assemblies, in order to minimize the false positive rate (<5%), maximize the N50 of the assembly and keep the virtual memory (95GB) and CPU (24 CPUs) requirements within a reasonable computational limit for our resources. In addition, we adjusted parameters at the mapping stage to create contigs, scaffolds and long-scaffolds to maximize N50 (overlap required in re-alignments, distance between mate-pairs, number of reads aligned to support assembly, see pipeline available at https://github.com/seb951/venustaconcha_ellipsiformis_genome).

Genome completeness was assessed using busco 3.0.2 (Benchmarking Universal Single-Copy Orthologs, Simao et al. 2015). Briefly, busco uses curated lists of known core single copy orthologs to produce evolutionarily-informed quantitative measures of genome completeness (Simao et al. 2015). Here, we tested both the eukaryotic (303 single copy orthologs) and metazoan (978 single copy orthologs) gene lists to assess the completeness of our genome assembly.

### Genome contamination

Mussels are filter feeders and tissues as gills can potentially harbor microbial fauna. In addition, freshwater mussels are prone to infection by trematodes (Müller et al. 2015, Capt et al. 2018). As suggested by Takeuchi *et al*. (2012), we first checked for the presence of double peaks in the distribution of the GC content of the raw reads, which would indicate contamination in the genome. In addition, we checked for potential contaminant sequences in the gene space of our current *V. ellipsiformis* genome assembly. Accordingly, we created a custom database of all trematodes, nematodes and bacterial protein sequences available from NCBI (39,617 and 156,174 protein sequences available from refseq database for trematodes and nematodes respectively, and 337,035 bacterial sequences available from uniprot database). We then compared this custom database of protein sequences to our predicted Open Reading Frames (315,932 *V. ellipsiformis* Open Reading Frames) to identify putative contaminants genes (blastp, mininum evalue of 1e-20 and minimum 90%/99% Percentage of identical matches).

### Characterization of repetitive elements

Given that repetitive elements can occupy large proportions of a genome, the characterization of their proportion and composition is an essential step during genome annotation. RepeatModeler open-1.0.10 (Smit & Hubley 2015) was used to create an annotated library of repetitive elements contained in the *V. ellipsiformis* genome assembly (excluding sequences <1Kb). Then, with RepeatMasker open-4.0.7 (Smit et al. 2015), we extracted libraries of repetitive elements for the taxa “Bivalvia” and “Mollusca” from the RepeatMasker combined database (comprising the databases Dfam_consensus-20170127 and RepBase-20170127) using built-in tools. Sequences classified as “artefact” were removed from the last two libraries before the subsequent steps. The three libraries were used alone and/or in combination (except for the Mollusca+Bivalvia combination) to mask the cut-down assembly again with RepeatMasker, specifying the following options: -nolow (to avoid masking low complexity sequences, which may enhance subsequent exon annotation), -gccalc (to calculate the overall GC percentage of the input assembly), -excln (to exclude runs of ≥20 Ns in the assembly sequences from the masking percentage calculations). Option -species was used to specify the taxon for the runs with Bivalvia and Mollusca libraries, while option -lib used to specify the *V. ellipsiformis* library and the combined ones. Results summaries for the latter three runs were refined with the RepeatMasker built-in tools. Linear model fit for genome size and repeats content for all available bivalve genomes were calculated with R, using the highest masking value found for *V. ellipsiformis*.

### Genome annotation

We used quast (Gurevich et al. 2013) to calculate summary statistics on the genome assembly. In addition quast uses GlimmerHMM (Majoros et al. 2004), a gene predictor that uses Hidden Markov Models to identify putative genes in the final assembly. Following this, we translated Open Reading Frames identified in the annotation files into protein sequences using bedtools v2.27.1 (Quinlan & Hall 2010) and the program transeq from emboss v6.6.0 (Rice et al. 2000) bioinformatics pipeline. These were then compared against the manually curated UniProt database (556,388 reference proteins, downloaded January 11^th^ 2018, e-value cut-off of 10^−5^) using BLASTp (Altschul et al. 1990). These steps were done on the long-scaffolds assembly, the masked long-scaffolds assembly (with low complexity regions replaced with N), in addition to the broken long-scaffolds assembly (scaffolds broken into smaller contigs by quast, based on long stretches of N nucleotides).

### Mitochondrial genome

Given the atypical mode of mitochondrial inheritance of freshwater mussels and therefore its evolutionary importance, we first aimed to check if the mitochondrial female genome had been properly assembled. Using BLASTn (Altschul et al. 1990) with high stringency (E value <1e-50), we identified a fragmented mitochondrial genome. We then created a mt specific dataset containing 1,396,004 sequence reads by aligning paired-end reads to the reference mt genome of Breton et al. (2009, GenBank Acc. No. FJ809753) using samtools v1.3.1 and bedtools v2.27.1 (Li et al. 2009; Quinlan & Hall 2010). We then rebuilt the mt genome *de novo* using abyss 2.0, testing different k-mers (17–45). In addition, we aligned reads to the reference transcriptome using bwa v0.7.12-r1039 (Li & Durbin 2009) and identified Single Nucleotide Polymorphisms (SNPs) with respect to the reference mt genome using samtools and bcftools v1.3.1 (Li et al. 2009).

## Results and Discussion

We generated 564M paired-end reads (2 × 100b) representing an average 65X coverage of the genome (**Table 1**). This was complemented by 98M mate-pairs (5Kb insert, 11X average genome coverage) and 103,000 Pacific Biosciences long reads (0.3X average genome coverage), and a recently published reference transcriptome comprised of 285,000 contigs (Capt et al. 2018). Filtering and trimming the raw paired-end and mate-pair sequences removed about 5% of the total base pairs from further analyses, indicating that the quality of the raw sequences was high (**Table 1**). K-mer analysis indicated that the number of unique k-mers peaked at 42. In addition, model fitting predicted a genome assembly size of 1.80 Gb (see **Figure 1** for k = 21, but note that similar values were found at other k-mer values analysed), which is smaller than the predicted genome size according to C-value for other freshwater mussel species in the order Unionida (*Elliptio* sp., c-value = 3; *Uniomerus* sp., c-value = 3.2), but in general agreement with the recent draft genome of other sequenced bivalves (0.55Gb-3.2Gb, see Table 2).

**Figure 1:**
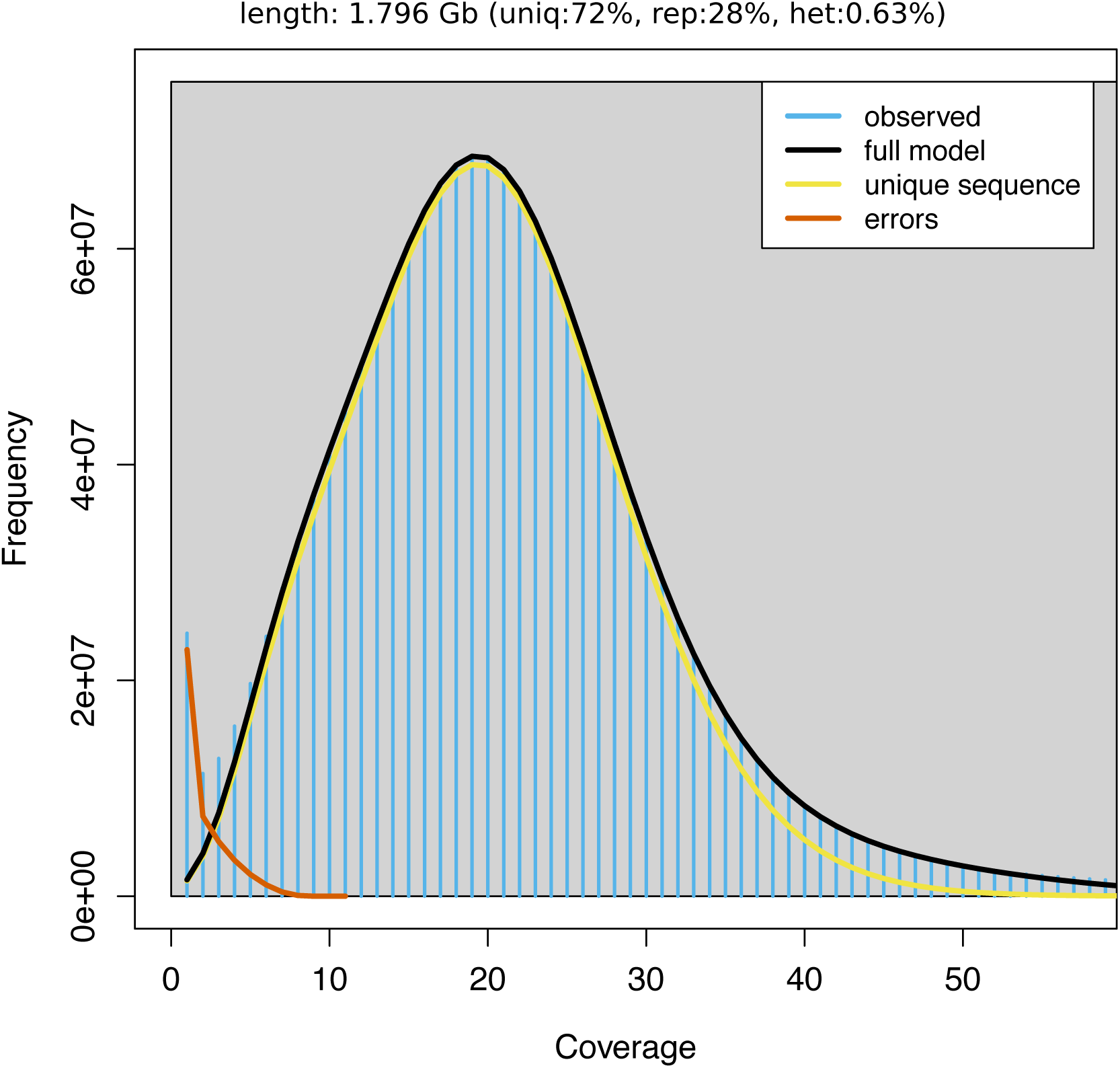
K-mer distribution (k=21) as calculated by genomescope (Vurture et al. 2017). Blue bars represent the observed k-mer distribution; black bar represent the modelled distribution without the error k-mers (red line) and up to a maximum k-mer coverage specified in the model (yellow line). Length: Estimated genome length, Uniq: Unique portion of the genome (non-repetitive elements), Rep: Repetitive portion of the genome, Het: Genome heterozygosity.

**Table 2:**
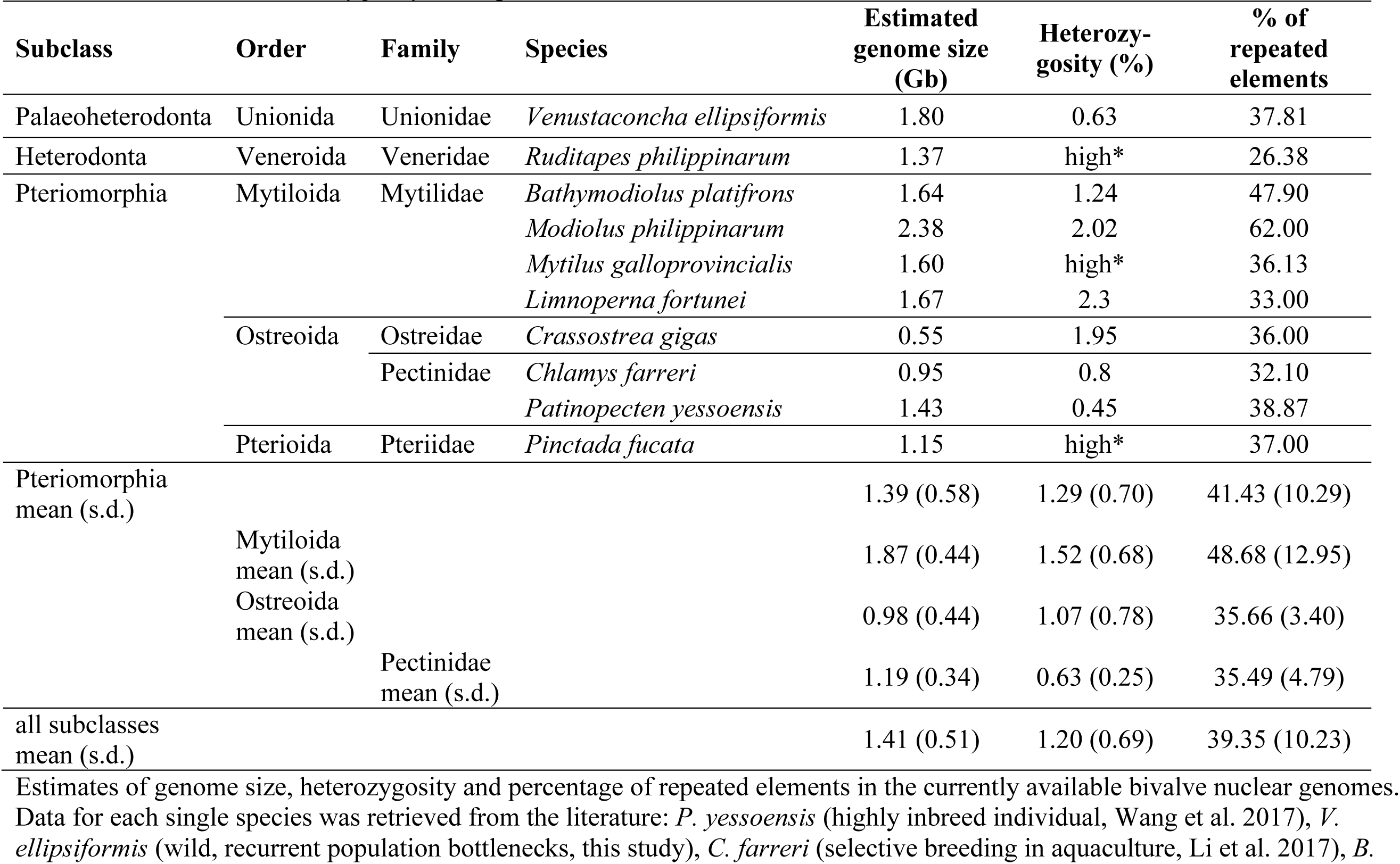
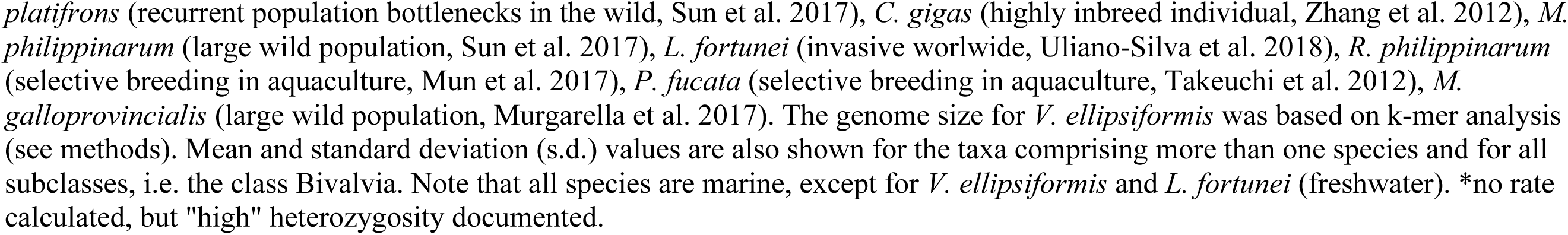
Genome size, heterozygosity and repeat elements

One of the main reason for the highly fragmentation of many bivalve genomes is thought to be high heterozygosity and repetitive elements. In fact, heterozygosity rates of most sequenced bivalve species appear to be high, even for highly inbred individuals under strong artificial selection for many generations (**Table 2**). In the current *V. ellipsiformis* genome assembly, we did not observe the typical double peak patterns in the k-mer distribution (see Figure 1) previously reported in most other bivalve genomes (Murgarella et al. 2016, Mun et al. 2016, Uliano-Silva et al. 2018). In fact, heterozygosity appears remarkably low (0.63%, Table 2) and more in line with previous reports for the deep-sea vent/seep mussel (Sun et al. 2017), where recurrent population bottlenecks as a result of population extinctions and re-colonizations of hydrothermal vents are common (Faure et al. 2015, Sun et al. 2017). Similarly, *V. ellipsiformis* populations have experienced severe genetic bottlenecks due to glaciation events (Zanatta & Harris 2012). The last glaciation in North America ended ~12,000 BP, after which individuals from glacial refuges were able to re-colonize previously uninhabitable regions. As a consequence, effective population size and heterozygosity for *V. ellipsiformis* is assumed to be fairly low, which was confirmed with the present dataset, and in contrast to most published bivalve genomes so far (Table 2).

Running the ABySS 2.0 assembly stage (abyss-bloom-dbg) led to a low False Positive Rate (<0.05%). The N50 for the contig assembly was 3.2Kb with 551,875 contigs (discarding contigs <1Kb, given that small contigs likely represent artefacts and provide little information for the overall genome assembly (Pavey et al. 2016; Murgarella et al. 2016, see **Table 3**). Once these were corrected and paired-end, mate-pairs and long read information were added, the scaffolds N50 increased to 5.5Kb, with 2.3% of nucleotides represented as “N” (see **Table 3** for the summary statistics and **Table 4** for overall genome assembly statistics acquired from quast analysis). Adding the Pacific Biosciences long reads only slightly improved the scaffolds N50 (from 5.5 to 5.7Kb, **Table 3**) and slightly decreased the number of *long-scaffolds* >1Kb (from 423,853 to 410,237), likely because our long read coverage was quite low (0.3X, **Table 1**). In addition, it is also possible that the more error prone Pacific Biosciences sequences, compared to Illumina paired-end reads, reduced their usability (Miller et al. 2017). Once the reference transcriptome was added, it improved the N50 to 6.5Kb, and substantially decreased the number of long-scaffolds to 366,926. This final long-scaffold assembly accounted for a total size of 1.54Gb (with 2.3% of “N” nucleotides) and represented 93% of the predicted genome size of 1.80Gb. Yet, it remained highly fragmented (366,926 scaffolds, **Table 3**). Genome annotation statistics can also be viewed in html format and downloaded here: https://github.com/seb951/venustaconcha_ellipsiformis_genome/tree/master/annotation_quast_v3

**Table 3:** Assembly statistics (ABySS2.0).

| assembly | n ( x10e6) | n:1000 | L50 | min | N80 | N50 | N20 | max | sum (Mb) |
| --- | --- | --- | --- | --- | --- | --- | --- | --- | --- |
| **raw unitigs** | 39.8 | 347,879 | 101,624 | 1,000 | 1,361 | 2,181 | 3,891 | 25,883 | 707 |
| **unitigs** | 18.5 | 444,734 | 127,617 | 1,000 | 1,485 | 2,452 | 4,273 | 25,944 | 984 |
| **contigs** | 14.0 | 551,875 | 141,012 | 1,000 | 1,704 | 3,117 | 5,817 | 39,408 | 1,449 |
| **scaffolds** | 13.7 | 423,853 | 92,607 | 1,000 | 2,303 | 5,477 | 9,099 | 45,260 | 1,539 |
| **long scaffolds (PacBio)** | 13.7 | 410,237 | 86,661 | 1,000 | 2,391 | 5,708 | 9,893 | 47,610 | 1,548 |
| **long scaffolds (PacBio + transcriptome)** | 13.6 | 366,926 | 58,906 | 1,000 | 2,534 | 6,523 | 16,660 | 298,135 | 1,549 |

n = number of contigs, n:1,000 = number of contigs of minimum length of 1,000, L50 = minimum number of sequences required to represent 50% of the entire assembly, min = minimum length of sequences analysed, N80, N50, N20 = weighted median statistic such that 80/50/20% of the entire assembly is contained in contigs equal to or larger than this value in bp, max = maximum size of contig in bp, sum = sum of all contigs of size > min, assembly stage (*raw unitigs* = raw assembly, not taking into account paired-end information, *unitigs* = filtering, merging and popping bubbles in *De Bruijn* graph, *contigs* = unitigs with paired-end information mapped, *scaffolds* = contigs with mate-pairs information mapped, *long scaffolds* = scaffolds with PacBio / transcriptome information integrated).

**Table 4:** Assembly and annotation statistics for the long scaffold assembly.

| **QUAST Assembly statistics** | **long_scaffolds** | **long_scaffolds (> 1 Kb scaffolds broken based on N streches)** | **long_scaffolds (> 1 Kb scaffolds, masked assembly)** |
| --- | --- | --- | --- |
| **Number of scaffolds ($\geq 0$ bp)** | 13,635,758 | 821,266 | 374,245 |
| **Number of scaffolds ($\geq 1$ Kb)** | 371,706 | 549,364 | 374,245 |
| **Number of scaffolds ($\geq 5$ Kb)** | 94,238 | 50,209 | 95,019 |
| **Number of scaffolds ($\geq 10$ Kb)** | 26,952 | 5,151 | 27,030 |
| **Number of scaffolds ($\geq 25$ Kb)** | 5,073 | 23 | 4,976 |
| **Number of scaffolds ($\geq 50$ Kb)** | 1,456 | 0 | 1,427 |
| **Total length ($\geq 0$ bp)** | 2,638,723,663 | 1,554,026,338 | 1,596,234,060 |
| **Total length ($\geq 1$ Kb)** | 1,590,292,198 | 1,425,294,273 | 1,596,234,060 |
| **Total length ($\geq 5$ Kb)** | 1,000,983,904 | 360,423,103 | 1,003,000,325 |
| **Total length ($\geq 10$ Kb)** | 541,545,133 | 64,766,821 | 538,648,016 |
| **Total length ($\geq 25$ Kb)** | 231,252,884 | 687,249 | 226,147,564 |
| **Total length ($\geq 50$ Kb)** | 107,178,666 | 0 | 104,739,660 |
| **Number of contigs** | 371,706 | 821,266 | 37,4245 |
| **Largest contig** | 313,274 | 44,597 | 31,3274 |
| **Total length** | 1,590,292,198 | 1,554,026,338 | 1,596,234,060 |
| **Estimated reference length** | 1,660,000,000 | 1,660,000,000 | 1,660,000,000 |
| **GC (%)** | 34.19 | 34.19 | 33.49 |
| **N50** | 6,656 | 2,812 | 6,627 |
| **number of N's per 100 kb** | 2,293.33 | 13.17 | 39,200.22 |
| **number of predicted genes (unique)** | 201,068 | 277,765 | 123,457 |
| **number of predicted genes ($\geq 300$ bP)** | 74,820 | 82,359 | 41,697 |
| **number of predicted genes ($\geq 1.5$ Kb)** | 18,539 | 14,338 | 11,897 |
| **number of predicted genes ($\geq 3$ Kb)** | 6,511 | 3,289 | 4,375 |
| **number of annotated ORF (uniprot)** | 29,031 | 14,198 | 25,544 |

All statistics are based on contigs of size $\geq 1$ Kb, unless otherwise noted (e.g., "# contigs ( $\geq 0$ bp)" and "Total length ( $\geq 0$ bp)" include all contigs.).

While assembly numbers (N50, number of scaffolds, etc.) are not directly comparable with other recently published genomes given the diversity of sequencing approaches (Illumina, 454, Sanger, PacBio), library types, sequencing depth and unique nature of the genome themselves, they can give a broad perspective of the inherent difficulties of assembling large genomes. The best comparison is probably with the saltwater mussel, *M. galloprovincialis*, giving their similar genome size (1.6Gb for *Mytilus* vs 1.80Gb for *V. ellipsiformis*) and Illumina paired-end sequencing approaches (32X for *Mytilus* vs 65X for *V. ellipsiformis*). While the *M. galloprovincialis* genome project (Murgarella et al. 2016) did not utilize mate-pair libraries or Pacific Bioscience long reads, they did make use of sequencing libraries with varying insert sizes (180, 500 and 800b). As such, they obtained a genome assembly quality relatively similar to ours and consisting of 393 thousand scaffolds (>1Kb), with however a substantially lower N50 (2.6Kb compared to 6.5Kb for *V. ellipsiformis*). The recently reported genome for the deep-sea vent/seep mussel *B. platifrons* (1.64Gb) made use of nine Illumina sequencing libraries with varying insert sizes (180 to 16Kb) and an overall coverage of >300X (Sun et al. 2017). With this very thorough sequencing approach and a heterozygosity closer to *V. ellipsiformis* than other Mytiloida, the scaffold N50 obtained was substantially higher (343.4Kb), but again the genome remained highly fragmented, into >65 thousands scaffolds. As exemplified here, high coverage sequencing libraries with varying insert sizes have become a broadly used approach for large and complex genomes. In fact, it is implemented by default in many genome assembly platforms (e.g. SOAPdenovo2, Luo et al. 2012; allpaths-lg, Gnerre et al. 2011). In the future, these libraries will likely be useful to further assemble the *V. ellipsiformis* genome, at least until these approaches are superseded by affordable, error free, single molecule long read sequencing (Gordon et al. 2016; Badouin 2017) or mapping approaches that allow reaching chromosome level assemblies such as optical mapping (e.g. Bionano Genomics, San Diego, CA).

Results of the busco (Simao et al. 2015) analyses showed that 664 (68%) of the 978 core metazoan genes (CEGs) were considered complete in our assembly. When the busco analysis was extended to include also fragmented matches, 871 (89%) proteins aligned. Results were similar when compared against the 303 core eukaryotic genes (61% complete, 86% complete or fragmented, **Table 5**). When compared to the previously published reference transcriptome for *V. ellipsiformis* (Capt *et al*. 2018), we found fewer complete genes, but also fewer duplicated genes (97.5% complete, and 24% duplicated in the reference transcriptome, compared to 68.1% complete and 1% duplicated here). This likely reflects the fact that the reference transcriptome is nearly complete, while the current reference genome is still fragmented. However, the reference transcriptome also likely contains multiple isoforms of the same genes, in addition to possible nematode contaminating sequences, despite the authors’ best efforts to minimize these problems. Previously analysed molluscan genomes of similar size (Murgarella et al. 2016; Sun et al. 2017) have found that 16% (*M. galloprovincialis*, 1.6Gb), 25% (pearl oyster *Pinctada fucata*, 1.15Gb), 36% (California sea hare *Aplysia californica*, 1.8Gb) of the core eukaryotic genes were complete. For their part Sun and collaborators (2017), identified 96% of the core metazoan genes to be partial or complete in the deep-sea vent/seep mussel *B. platifrons* (1.6Gb), again reflecting that the depth and type of sequencing, in addition to the idiosyncrasies of each genome, can have considerable influence on the end results.

**Table 5:** Analysis of genome completeness using BUSCO 3.0.2 (Benchmarking Universal Single-Copy Orthologs, Simao et al. 2015).

|  | metazoa | eukaryota |
| --- | --- | --- |
| Complete orthologs (C) | 664 (68%) | 185 (61%) |
| Complete and single-copy orthologs (S) | 652 (67%) | 181 (60%) |
| Complete and duplicated orthologs (D) | 12 (1%) | 4 (1%) |
| Fragmented orthologs (F) | 207 (21%) | 76 (25%) |
| Missing orthologs (M) | 107 (11%) | 42 (14%) |
| Total ortholog groups searched | 978 | 303 |

We confirmed the presence of a single GC content peak (Figure S1), thus supporting a lack of sequence contamination in the raw paired sequencing reads. In addition, we identified a very small percentage of Open Reading Frames matching to our custom database of nematodes-trematodes-bacteria proteins. Out of 315,932 Open Reading Frames identified in *V. ellipsiformis*, we identify 299 and 29 proteins with greater than 90% and 99% sequence identity (between 0.09% and 0.0009% of all ORF, respectively, Supplementary Table S1). This confirms that in the current genome assembly, the gene space is effectively free of the most common contaminants of freshwater mussels.

The custom *V. ellipsiformis* repeat library created *de novo* with RepeatModeler contained 2,068 families, the majority of them (1,498, 72.44% of the total) classified as “unknown”. Repeat content values reported below are slightly higher than the ones calculated based on k-mer analyses (28%, Figure 1), but should be considered more accurate given that they are based on the assembled sequences, rather than raw reads. The genome masking performed with the Bivalvia and Mollusca libraries had scarce performances (masking 2.38% and 2.59%, respectively; details in **Supplementary Table 2**), possibly because of the phylogenetic distance between *V. ellipsiformis*, which belongs to the early-branching bivalve lineage of Palaeoheterodonta, and the other bivalve and mollusk species represented in the database as well as their relative number of sequences. The custom *V. ellipsiformis* library masked 37.17% of the genome, while the combined *V. ellipsiformis* +Bivalvia masked 37.69% of the genome and the *V. ellipsiformis* + Mollusca reached 37.81%, the highest masking percentage (**Supplementary Table 3**). After refining, these raw values slightly decreased to respectively 36.29%, 36.80%, and 36.91% (**Supplementary Table 4**). All these latter values of repeat content fall in the 32–39% range (the median for all species is 37%) where six out of the nine sequenced bivalve species lie, irrespective of their genome size (*M. philippinarum* and *R. philippinarum* are the furthest from this interval) (**Table 2** and **Supplementary Figure 2**). Although the number of species sequenced up to now is still low, this observation indicates that repetitive elements may contribute differently to the total genome size among the different bivalve taxa: indeed, the correlation between genome size and repeats content is weak (**Supplementary Figure 2**). In both the *ab initio* masking with the *V. ellipsiformis* library and the two combined ones, most of the identified repeats are categorized as “unknown” (22.8% of the assembly), followed by retroelements (LINEs 2.9%, LTR elements 2.3–2.4%, and SINEs 1.7%, for a total of 6.9% of the assembly) and DNA elements (5.4–5.6% of the assembly) (**Supplementary Table 4**). Direct comparisons of these values with other species should be performed with caution, as the usually large “unclassified” portion of repeats might contain species-specific variants of known elements (Murgarella et al. 2016) that may therefore change the relative weight of each category on the total.

quast was used to calculate summary statistics and identify putative genes in the final assembly using a hidden markov model (**Table 4**). Following this, 29,031; 14,195 and 25,544 Open Reading Frames were annotated using BLASTp against UniProt database in the long-scaffolds, broken and masked long-scaffolds assemblies, respectively.

Freshwater mussels, marine mussels, as well as marine clams are the only known exception in the animal kingdom with respect to the maternal inheritance of mitochondrial DNA (see Breton et al. 2007 for a review). Their unique system, characterized by the presence of two gender-associated mitochondrial DNA lineages, has therefore attracted studies to better understand mitochondrial inheritance and the evolution of mtDNA in general. Using blastn, we recovered 53 contigs matching to the 15,975bp female reference mt genome from Breton *et al*. (2009), indicating that the mt genome was highly fragmented and likely improperly assembled with our current approach, much like what was found in the *M. galloprovincialis* genome draft of Murgarella et al. (2016). As such, we created a dataset of mt specific sequences that could be aligned to the mt genome (1,396,004 reads). This mt specific dataset was then re-assembled *de novo*, using different k-mers (17–45). Using a k-mer similar or larger to the one used in the overall assembly (k≥41) resulted in a failed assembly (no contigs created, data not shown), while using a k-mer <21 generated a highly fragmented mt genome (data not shown). Using a k-mer between 21 and 39 generated one large contig of 16,024bp comprising the entire mitogenome, with a 42bp insertion in the 16S ribosomal RNA. Given the different rate of evolution of mtDNAs, it is likely that assembly parameters we used for the whole genome were not appropriate for the *V. ellipsiformis* female mt genome. Finally, we also re-aligned the mt specific dataset to the original mt genome of Breton et al. (2009) and found high coverage (mean = 7,256X, SD = 682) for most positions, while for three regions coverage dropped below 300X (**Figure 2**). Six SNPs with respect to the reference were also identified, indicating possible polymorphism, or sequencing error in the original mt reference genome (**Figure 2**).

**Figure 2:**
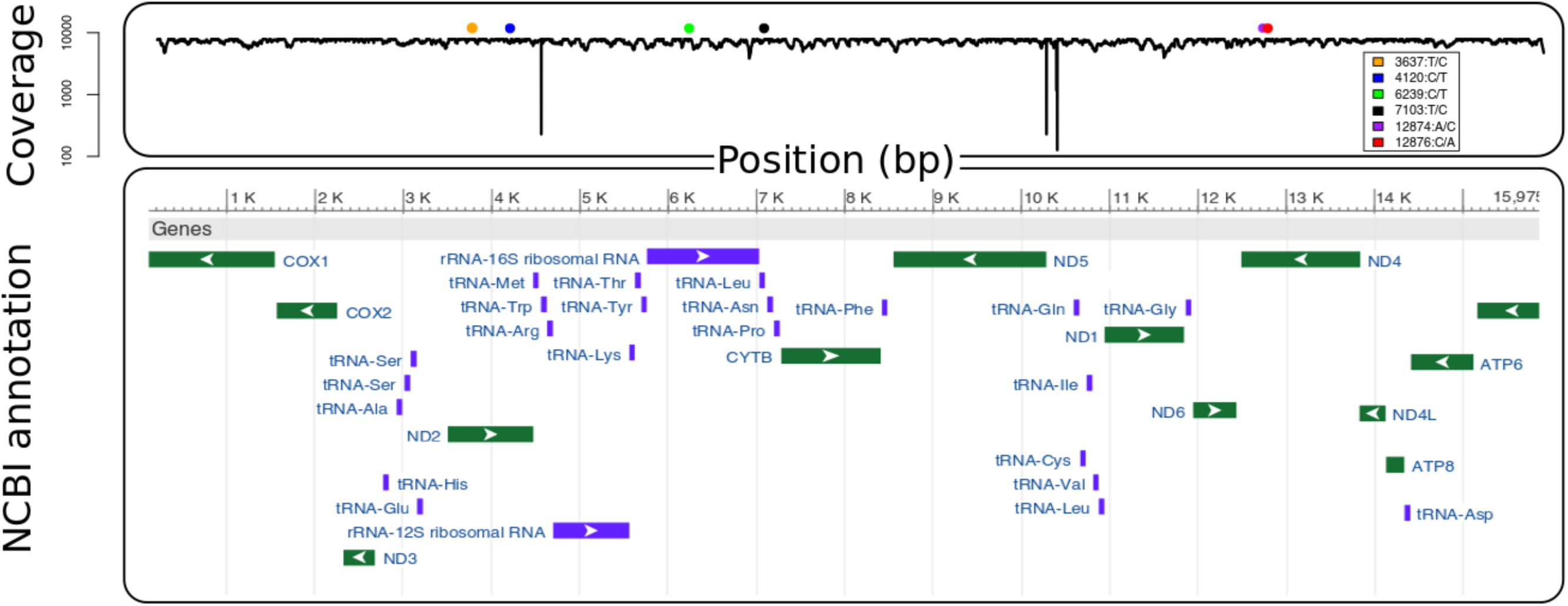
Mitochondrial coverage based on sequence alignment and annotation (from NCBI). Six nucleotide positions were identified in the legend as fixed for an alternative allele compared to the reference of Breton et al. (2009).

## Conclusion

High throughput sequencing has the power to produce draft genomes that were only reserved to model systems ten years ago. Here we report the first *de novo* draft assembly of the *V. ellipsiformis* genome, a freshwater mussel from the bivalve order Unionida. Our assembly covers over 86% of the genome and contains nearly 90% of the core eukaryotic orthologs, indicating that it is nearly complete. In addition, we calculated relatively low heterozygosity rates, uncommon in bivalves, but likely explained by the recent evolutionary history of *V. ellipsiformis*. Finally, as for other mussel genomes recently published, our genome remains fragmented, showing the limits of high throughput sequencing and the necessity to combine different sequencing approaches to augment the scaffolding and overall genome quality, especially when a large fraction of the genome is comprised of repetitive elements. In the future, the *V. ellipsiformis* genome will benefit from a larger number of long read sequences, varying library size for paired-end sequencing, and the use of genetic, physical or optimal maps to subsequently order scaffolded contigs into pseudomolecules or chromosomes.

**Abbreviations**
BLAST: Basic Local Alignment Search Tool
bp: base pairs
CPU: Central Processing Unit
DNA: Deoxyribonucleic acid
Gb: Gigabases
GB: gigabytes
Kb: Kilobases
LINEs: Long interspersed elements
LTR: Long terminal repeats
L50 =: minimum number of sequences required to represent 50% of the entire assembly
M: Million
Mb: Megabases
mt: mitochondria
N80/50/20: weighted median statistic such that 80/50/20% of the entire assembly is contained in contigs/scaffolds equal to or larger than this value.
ORF: Open Reading Frames
RAM: Random Access Memory
SINEs: Short interspersed elements

## Data availability

Supporting data for this Genome Report will be made available on datadryad.org upon acceptance Raw sequences are available in the SRA database with number SRP132483 (submission SUB3624229 to be release upon publication) and Bioproject accession PRJNA433387. All scripts used in the analyses are available on github (https://github.com/seb951/venustaconcha_ellipsiformis_genome).

## Acknowledgments

Computations were made on the supercomputer briaree from Université de Montréal, managed by Calcul Québec and Compute Canada. The operation of this supercomputer is funded by the Canada Foundation for Innovation (CFI), the ministère de l’Économie, de la science et de l’innovation du Québec (MESI) and the Fonds de recherche du Québec - Nature et technologies (FRQ-NT).

